# The effects of *de novo* mutation on gene expression and the consequences for fitness in *Chlamydomonas reinhardtii*

**DOI:** 10.1101/2022.09.06.506807

**Authors:** Eniolaye J. Balogun, Rob W. Ness

## Abstract

Mutation is the ultimate source of genetic variation, the bedrock of evolution. Yet, predicting the consequences of new mutations remains a challenge in biology. Gene expression provides a potential link between a genotype and its phenotype. But the variation in gene expression created by *de novo* mutation and the fitness consequences of mutational changes to expression remain relatively unexplored. Here, we investigate the effects of >2600 *de novo* mutations on gene expression across the transcriptome of 28 mutation accumulation lines derived from two independent wild-type genotypes of the green algae *Chlamydomonas reinhardtii*. We observed that the amount of genetic variance in gene expression created by mutation (*V_m_*) was similar to the variance that mutation generates in typical polygenic phenotypic traits and approximately 15-fold the variance seen in the limited species where *V_m_* in gene expression has been estimated. Despite the clear effect of mutation on expression, we did not observe a simple additive effect of mutation on expression change, with no linear correlation between the total expression change and mutation count of individual MA lines. We therefore inferred the distribution of expression effects of new mutations to connect the number of mutations to the number of differentially expressed genes (DEGs). Our inferred DEE is highly L-shaped with 95% of mutations causing 0-1 DEG while the remaining 5% are spread over a long tail of large effect mutations that cause multiple genes to change expression. The distribution is consistent with many *cis*-acting mutation targets that affect the expression of only one gene and a large target of *trans*-acting targets that have the potential to affect tens or hundreds of genes. Further evidence for *cis*-acting mutations can be seen in the overabundance of mutations in or near differentially expressed genes. Supporting evidence for *trans*-acting mutations comes from a 15:1 ratio of DEGs to mutations and the clusters of DEGs in the co-expression network, indicative of shared regulatory architecture. Lastly, we show that there is a negative correlation with the extent of expression divergence from the ancestor and fitness, providing direct evidence of the deleterious effects of perturbing gene expression.

## Introduction

Mutations provide both the raw variation for evolution through adaptation and contribute towards disease and aging. Unfortunately, a challenge in predicting the effects of mutations is linking the genotypic consequences of a mutation to phenotypic change. Gene expression provides a potentially useful link where we can observe changes in the regulation of genes due to mutation. Moreover, there is a longstanding hypothesis that evolution may proceed primarily through alteration in gene expression rather than changes in protein structure (King and Wilson 1975). For this to be true, mutations altering expression must either be more common, have less harmful effects or be more beneficial than protein-coding mutations. Though there are now numerous examples of regulatory changes driving both adaptive changes (e.g. Abzhanov et al. 2006) and harmful diseases (e.g. Kleinjan and van Heyningen 2005), we still lack a general description of how variation in gene expression originates from *de novo* mutation and the distribution of effects of these mutations on fitness.

Comparative research has revealed numerous patterns in gene expression variation that provide insights into the impact of mutation on gene regulation (reviewed in Hill et al. 2021). In general, essential genes, genes typically with high expression and those that are central in molecular interaction networks tend to display reduced expression variation within and between species (Jeong et al. 2001; Pál et al. 2001; Yu et al. 2004; Hahn and Kern 2005; He and Zhang 2006; Liao et al. 2006; Yu et al. 2007; Zotenko et al. 2008). An obvious explanation is that stabilizing selection on expression level removes deleterious variants and reduces the overall segregating expression variation. However, the distribution of mutational effects on expression (DEE) may also influence the extent of variation in different gene classes if mechanisms have evolved that buffer essential genes from perturbations by mutation. It has been observed that within species variation is driven by *trans*-regulatory variants, defined by regulatory mutations that affect distal genes. In contrast, *cis*-regulatory mutations, which affect proximal genes, underlie between-species variation (Wittkopp et al. 2004; Wittkopp et al. 2008; Graze et al. 2009; Signor and Nuzhdin 2018). This trend might be explained if *trans*-regulatory mutations are more common, due to a larger mutation target. The more genes affected, the increasing likelihood of reducing fitness (Zande et al. 2022). With this in mind, we may also expect *trans*-regulatory variants to be kept at low frequencies and thus not contribute to between species divergence. Critically, the above explanations all depend on the rate, target size, expression effect and fitness impact of regulatory mutations, highlighting the importance of studying the effect of mutation on expression.

Some of the most detailed information on how *de novo* mutation affects gene expression comes from experiments using single gene models where the effect of mutations on a reporter gene can be studied in detail. For example, in *S. cerevisiae, cis*-acting mutations in the *TDH3* promoter were shown to generally reduce expression and have a greater effect on expression than *trans*-acting mutations, which did not show an asymmetry in the direction of effect (Metzger et al. 2016). However, although *trans*-acting mutations were relatively weak, they were 265 times more likely to occur than *cis*-acting mutations, implying a much larger mutational target (Gruber et al. 2012; Metzger et al. 2016). Despite the valuable insights gained from such precise experiments, these systems can not capture the diverse mechanisms that may govern the effects of natural mutations on expression throughout the transcriptome. For example, some genes have redundant promoters and regulatory proteins that maintain expression in the presence of mutational perturbations (reviewed by Payne and Wagner 2015; Signor and Nuzhdin 2018). Network feedback may also serve to mute the effects of mutation on expression of genes, but such effects would likely be most evident when genes native to the genome in question are investigated. Additionally, there is evidence that *trans*-acting eQTLs may be clustered in hotspots, affecting thousands of genes (Albert et al. 2018), which are not observable in single gene model systems. There is therefore good reason to investigate the effect of *de novo* mutation on the transcriptome.

Investigating spontaneous mutations has been constrained by the rarity with which new mutations arise. Mutation accumulation (MA) experiments overcome this limitation by evolving lines under minimal selection, therefore facilitating the build-up of spontaneous changes irrespective of their fitness (Muller 1928). MA has been used extensively in the estimation of the rate, spectrum, and fitness effects of mutations. However, there are surprisingly few studies investigating the expression changes in MA lines in a small number of model organisms including *S. cerevisiae*, *D. melanogaster* and *C. elegans* (Denver et al. 2005; Rifkin et al. 2005; Landry et al. 2007; McGuigan et al. 2014; Huang et al. 2016; Zalts and Yanai 2017). A key parameter in measuring how mutation affects gene expression is the contribution of mutation to expression variation per generation *i.e.,* mutational variance (*V_m_*) (Lynch and Walsh 1998). From the studies formerly mentioned, the mutational variance of *C. elegans* (Denver et al. 2005) was up to 9-fold higher than that of *D. melanogaster* and (Rifkin et al. 2005) and yeast (Landry et al. 2007). MA studies have also shown that mutation introduced far more variation in expression than would be expected based on standing expression variation in natural populations, suggesting strong stabilizing selection in nature (Lemos et al. 2004; Denver et al. 2005; Rifkin et al. 2005; Huang et al. 2016). We have also learned that the expression of genes likely involved in fitness-related traits and pivotal developmental stages are highly constrained (Zhang and Li 2004; Rifkin et al. 2005; Zalts and Yanai 2017). Moreover, there is evidence of an inverse correlation between *V_m_* and expression level (Rifkin et al. 2005). Despite these breakthroughs (Hine et al. 2018; Hodgins-Davis et al. 2019), our current knowledge of the global effects of mutations across the transcriptome is restricted to a small number of organisms with sometimes limited technologies and often in the absence of information about the underlying genomic mutations.

Here, we investigate the effect of *de novo* mutation on gene expression and fitness in *Chlamydomonas reinhardtii*. This haploid green alga is a longstanding model for numerous aspects of cell, molecular, and genome biology (Harris 2001; Grossman et al. 2003) and has emerged as a model for studying the evolution of mutation in eukaryotes. Over the past decade, numerous studies have explored mutation rate (Ness et al. 2012; Sung et al. 2012; Ness et al. 2015; Ness et al. 2016; Flynn et al. 2018), base spectrum, and the predictors of mutational variation across the genome of *C. reinhardtii* and related species (Ness et al. 2015; López-Cortegano et al. 2021). We also know that on average mutations in *C. reinhardtii* are deleterious, but evidence from the genetic mapping of mutational effects demonstrates a significant fraction of spontaneous mutations are beneficial (Morgan et al. 2014; Böndel et al. 2019). However, in this system, it remains unexplored how the accumulation of mutations has altered patterns of gene expression. Here, we sequenced the transcriptomes of 28 well-studied MA lines derived from two independent wild-type genotypes with fully characterized mutations and known fitness (Morgan et al. 2014; Ness et al. 2015; Böndel et al. 2019). Each MA line was grown for 800 – 1100 generations and carries approx. 76 mutations. Integrating new transcriptome data with detailed characterization of these lines, we address the following questions: (1) How much does *de novo* mutation influence the direction and overall variation in gene expression? (2) What is the distribution of mutational effects on expression? (3) What are the properties of genes that are more or less robust to gene regulation perturbation by mutation (4) Do MA lines with more transcriptome divergence from mutation also suffer greater fitness declines?

## Methods

### Sample collection and growth

We examined gene expression in 28 mutation accumulation lines derived from two wild-type *C. reinhardtii* strains: CC-2391 (n=13) and CC-2344 (n=15) (Morgan et al. 2014; Ness et al. 2015; Böndel et al. 2019). These MA lines have been intensively studied, including the characterization of the location of each mutation; the relative fitness of each MA line with respect to their unmutated ancestral line; the mutation rate of each line; and the distribution of fitness effects of all mutations. The MA lines were grown for 800-1100 generations and carry a mean of 76.2 mutations in CC-2344 and 112.1 mutations in CC-2931. To measure expression change from *de novo* mutation, we grew three replicates of each MA line and their unmutated ancestors synchronously on Bold’s agar medium at 25°C under standard, uniform lab conditions. Each replicate was grown until the density of colonies was sufficient for RNA isolation, then frozen at −80°C until extraction. RNA extraction was conducted via the Maxwell RSC 48 Instrument and assessed for degradation, purity, and quantity using gel electrophoresis and a Qubit fluorometer. mRNA isolation and library preparation were conducted with the NEB mRNA stranded library preparation kit and Illumina NovaSeq preparation. All samples were sequenced using Illumina 100bp paired-end sequencing with at least 20 M high-quality reads per sample. Sequencing and library preparation were performed by Genome Quebec.

### Sequence read filtering and quality control

Sequence quality of raw reads was assessed with FastQC (*v0.11.9*) (Andrews 2010) and aligned using STAR aligner (*v2.7.8a*) (Dobin et al. 2013). We used the *C. reinhardtii* v6.0 reference genome with the mt+ mating type allele, and organelle genomes included (Craig et al. 2022). STAR alignment was conducted in a two-pass mode, where the first pass identified possible splice junctions (SJ), and the second pass used the reference gene annotation in conjunction with the SJ output. Parameters of both alignments included “--alignIntronMax 5000” to exclude known false large introns identified in other STAR-mapped BAM files and “--outFilterMismatchNoverLmax 0.2”, which allowed 0.2 x 100 bp mismatches for each single read to account for *C. reinhardtii’s* high genetic diversity (∼3%). The resulting BAM files were sorted and indexed with samtools (Li et al. 2009). Sequence coverage and mapping quality was measured with bamqc and rnaseq from the Qualimap quality control package (*v2.2.2a*) (Okonechnikov et al. 2016). Lastly, transcript quantification was done with featurecounts (*v2.0.0*) (Liao et al. 2013).

### Differentially expressed gene (DEG) calling

Read count normalization was conducted by DESeq2 (*v1.30.1*) (Love et al. 2014) using the median of ratios method to account for differing sequencing depth across samples. Low read counts were filtered by DESeq2 using the mean of normalized read counts across all samples within a strain (*i.e.,* ancestors and replicates) as a filter statistic that optimizes statistically significant differentially expressed gene calling. We retained ∼ 95% of all genes across samples. We found a high expression correlation between the biological replicates of all samples using FPKM (fragments per kilobase of exon per million) normalized read counts of each gene (Pearson’s *R* > 0.96, *p* < 1.0 x 10^-4^). Of the 90 replicates, only two had a lower correlation with their respective replicates (CC-2344 L6 replicate 2, Pearson’s *R* ∼ 0.78, *p* < 1.0 x 10^-4^; CC-2931 L3 replicate 3, Pearson’s *R* ∼ 0.80, *p* < 1.0 x 10^-4^). These replicates were removed from all analysis dependent on averaged normalized read counts, but all replicates were kept for calling differentially expressed genes (DEGs) given DESeq2 accounts for variance among replicates. DEGs in each MA line were identified with respect to its unmutated ancestral line with DESeq2. DESeq2 uses the Wald’s test, *p*_adj_ < 0.05 for DEG calling, and the Benjamini-Hochberg method to account for multiple testing (*FDR < 0.05*). Across all MA lines, we identified 23,557 DEGs in CC-2344 (mean ∼1571) and 15,829 DEGs in CCC-2391 (mean ∼1218).

### Quantifying expression variance introduced by de novo mutation

To estimate the mutational variance, we used a generalized linear mixed model with a Poisson distribution to find the between line variation across MA lines.

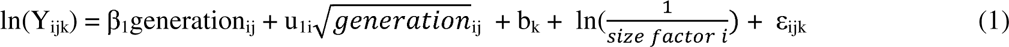

The model was fitted for each gene separately, where the between line variance per generation is given by the variance of u_1i_ (2Var(u_1i_) ∼ mutational variance, *V_m_*) (Lynch and Walsh 1998). The MA line was treated as a random effect u_1i_ ∼ N(0,σ^2^_u_) to account for per line variation given the MA lines evolved independent of each other. Since we know the total number of mutations in each line, we were able to estimate the increase in expression variation per *de novo* mutation (*V_mm_*) by substituting the number of generations with the mutation count in equation 1. In our model, Y_ijk_ is the raw read count of gene j, line i and replicate k, which is normalized by a size factor 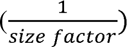 to consider the varying sequencing depths across samples. Though we found a strong correlation between replicates, we also included an observation level random effect b_k_ ∼ N(0,σ^2^_b_) to account for overdispersion in our data. To estimate the mutational heritability 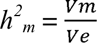, the environmental variance (*V_e_*) for each gene is given by the residual variance of the random slope ∼ σ^2^_ε_. For subsequent analysis, all variance estimates were log_2_ transformed to facilitate comparison with previous studies.

To test whether lines with more mutations demonstrated larger expression divergence from the unmutated ancestor, we quantified the log_2_-fold change of gene expression for each gene in each MA line relative to its respective unmutated ancestor. We then summed these expression changes for all significant DEGs (*p*_adj_ < 0.05) for each MA line and plotted it against the mutation count of the respective line to get an overall measure of divergence (Fig 1). The DEGs in each MA line were divided into positive and negative expression change bins to consider any bias in the direction of change and avoid the cancelling out of expression changes. We found no correlation between expression divergence and mutation count and found similar results after plotting the sum of absolute fold change of DEGs per MA line against mutation count (Fig S1).

**Fig 1.**
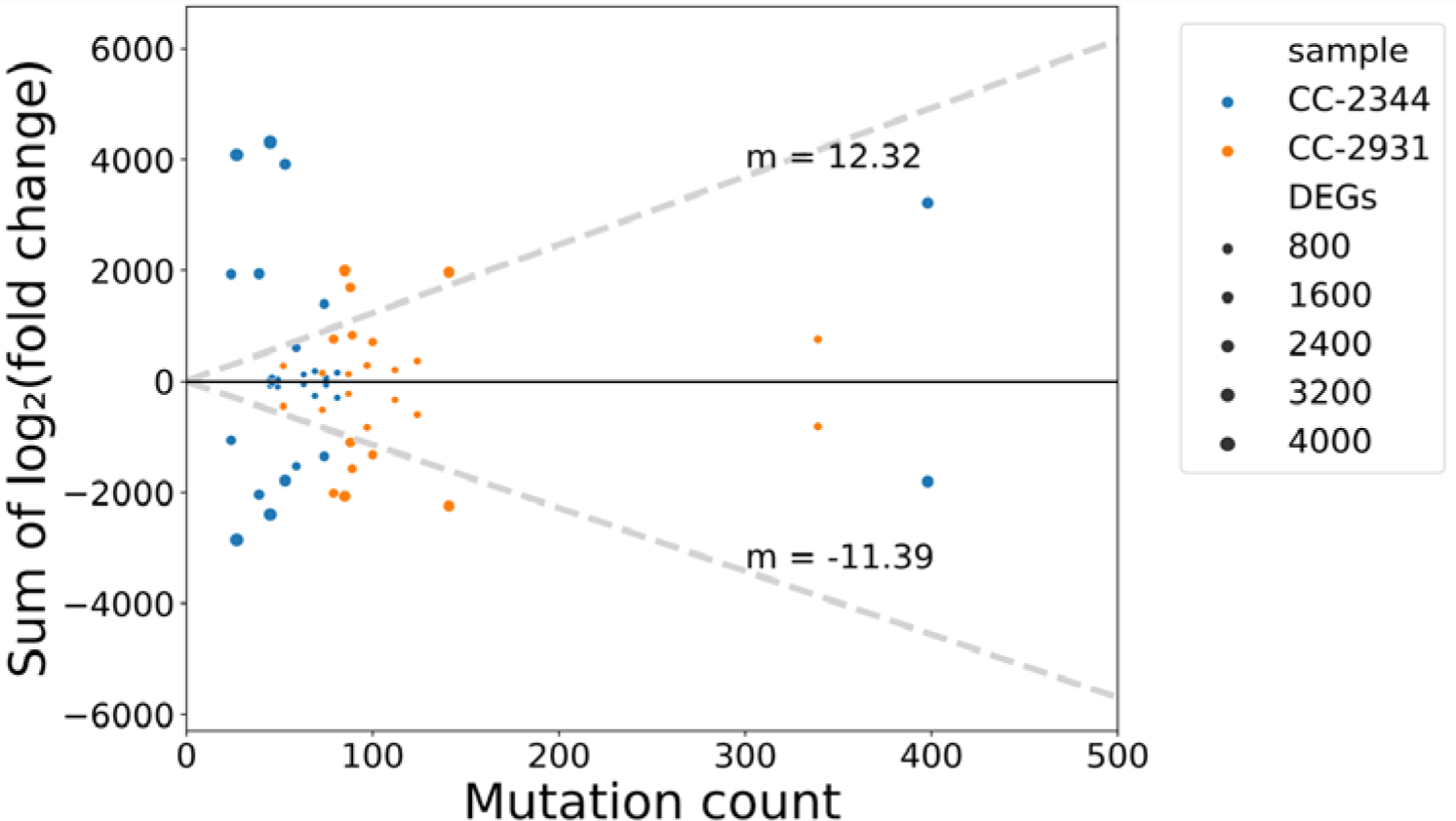
No correlation between total expression change and mutation count. Plot of the total expression change of differentially expressed genes in each MA line against mutation count. Genes with positive and negative log_2_ fold changes were summed separately and their total expression change was represented as points above and below the black horizontal line. The dashed line represents the average directional rate of 2-fold expression change per mutation. There is no correlation between expression change in either direction and the number of mutations (positive: Pearson’s *R* = 0.1, *p* = 0.625; negative: Pearson’s *R* = 0.08, *p* = 0.702).

### Inference of the distribution of expression effects (DEE) of novel mutations

The number of DEGs generated by mutation is a function of the distribution of expression effects of new mutations. Given that many mutations likely have no effect on expression, while other mutations may affect many genes (reviewed in Emerson and Li 2010), we inferred the distribution of expression effects of mutations (DEE) by fitting a model to the number of DEGs caused by mutation in each of the 28 MA lines. We first fit a model where the number of DEGs caused by a mutation was drawn from a gamma distribution:

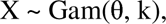

where X is the number of DEGs and θ and k are the parameters for the shape and scale respectively. We also reasoned that if many mutations caused 0 DEGs a better fitting model may allow a fraction of mutations to cause 0 DEGs while other mutations caused 1 or more DEGs drawn from a gamma distribution shifted by 1. To find the parameters that best optimize the likelihood of seeing all 28 mutations-DEG counts in a given distribution, we use the SciPy global minima tool dual annealing, which was amended to find the global maximum instead of the minimum (Virtanen et al. 2020). The tool was run for one million iterations under varying temperatures to get a good estimate of the optimum. For the modified distribution, we split the bounds of no-effect mutations into 0.2 intervals and ran the model for each interval. The model of best fit was determined through Akaike and Bayesian Information criteria.

### Correlates of expression change

To test whether expression change depends on transcription level in the ancestral genotype, we categorized the genes into 10 percentile bins according to the FPKM (fragments per kilobase of exon per million)-normalized read counts of the ancestral line. The lowest bin represents genes with the lowest read counts and those in the highest percentile bins consist of high expression genes. Each bin contains 1745 and 1736 genes for CC-2344 and CC-2931 respectively. Next, we plotted the distribution of expression change within each percentile bin by taking the median log_2_ fold change of each gene across all MA lines within a strain.

Using genes in the metabolic network (iCre1355), we tested whether the position of genes in the network, specifically their connectedness to other genes predicted their robustness to mutational perturbations. The degree centrality (*i.e.,* indegree, outdegree and betweenness centrality) of each gene was estimated (Imam et al. 2015) from the iCre1355 network after conversion to a gene centric network (Chaiboonchoe et al. 2016). iCre1355 was constructed using the *C. reinhardtii* v5 gene annotation with 1355 metabolic genes, and after liftover to the v6 annotation, 1304 metabolic genes remained.

We also tested whether genes central in the co-expression network were buffered against expression change due to mutation. We identified hub genes in the Strenkert et al (2019) co-expression network that measured the expression of the transcriptome at two hour intervals over a 24-hr period. Using the log_2_(FPKM + 1) normalized expression data retrieved from the Strenkert et al. (2019) supplementary, we generated an adjacency matrix with a soft threshold of 2 using WGCNA (Langfelder and Horvath 2008; Langfelder and Horvath 2012). The adjacency matrix was subsequently transformed into a topological overlap matrix which was used to cluster coexpressed genes using hierarchical clustering analysis. Of the 33 resultant clusters, those that were correlated by 0.75 were merged to provide a total of 13 distinct clusters. 99.9% of genes were placed within these 13 co-expression modules. Hub genes were identified as genes with the top 5% intramodular connectivity per module. There were 890 hub genes based on the v5 gene annotation, which translated to 774 hub genes in v6. We measured the relationship between expression change and intramodular/total connectivity using multiple regression and compared the median expression change in hub genes against non hub genes.

### Evidence of cis- and trans-acting mutations

To identify putative *cis* mutations, we measured the number of mutations located within 100 bp of the nearest DEG. Given the possibility of random chance neighboring mutation-DEG pairs, we use a permutation test to identify the presence of significant mutation-DEG clustering. We compared the number of nearby mutations to *n* observed DEGs per MA line to a null distribution generated by 10,000 iterations of nearby mutations to *n* randomly selected DEGs. The *p* value, the fraction of simulated DEGs with a higher number of mutation-DEG clustering than our real dataset, was subsequently compared to α = 0.05 to test for the presence of potential *cis* mutations.

To group DEGs that might arise from mutations in common regulatory elements, we considered whether DEGs were clustered in the *Chlamydomonas* co-expression network (Strenkert et al. 2019). To test for excess clustering, we calculated the shortest path in a network between DEGs in the same MA line using the multi_source_dijkstra function in the NetworkX python package (Hagberg et al. 2008). Paths between genes (*i.e.,* edges) in the co-expression network represent correlations in gene expression. Since genes can be connected through hubs, we measured total connectedness by converting each correlation coefficient (weight of the path) to path length using the equation: distance = correlation coefficient^-1^. We then took the sum of all paths between two genes to get the total distance between two DEGs in the coexpression network. The shortest total paths represent the strongest correlations. We used a permutation test to compare the distribution of shortest path lengths between observed DEGs for each MA line against a null distribution generated from the shortest path lengths of *n* random genes, where *n* is the number of DEGs in a given MA line. Lines with signs of significant clustering had shorter distances than their simulated counterparts. A *p* value was assigned as the fraction of trials where simulated DEGs had equal or shorter path lengths than the observed data. The function multi_source_dijkstra from NetworkX was amended to exclude unconnected DEGs from the analysis.

### Fitness of mutation accumulation lines

The fitness of all 28 MA lines were previously characterized in Morgan et al (2014). Fitness was measured as the mean of maximum growth rate from two independent assays in Bold’s medium. To assess fitness change, all fitness measures were scaled relative to the growth rate of the unmutated ancestor. To test for an association of fitness change with expression change, we performed a linear regression of relative fitness against the number of DEGs per line. We performed a similar regression using absolute fold expression change which resulted in qualitatively similar results (Fig S6).

The code used for the analysis can be found at https://github.com/enibalogun/The-effect-of-mutation-on-gene-expression.git and the raw data at the European Genome-Phenome Archive [PRJEB51708/ERP136361].

## Results and Discussion

### Mutational variance

In this study, we examined how >2600 mutations across 28 MA lines have altered gene expression in the transcriptome of *C. reinhardtii*. On average each MA line had 1407 differentially expressed genes (DEGs), with more DEGs in MA lines of strain CC-2344 (mean 1571 DEGs/MA line) than CC-2931 (mean 1218 DEGs/MA line). We first estimated mutational variance (*V_m_*) (Lynch and Walsh 1998), which describes the change in expression variation introduced into each gene in each generation due to mutation. Measured as a log_2_-fold change in RNASeq read count, we found that the median *V_m_* across genes was 8.2 x 10^-4^ and 6.1 x 10^-4^ for CC-2344 and CC-2931 respectively. Similar to the mean number of DEGs, the *V_m_* per gene was higher in MA lines derived from CC-2344. The 1.3-fold elevation in CC-2344 was unexpected given that it has a lower mutation rate than CC-2931 (Ness et al. 2015). Our estimates of *V_m_* for gene expression were on average 1.8 X that of *C. elegans* (Denver et al. 2005; Hodgins-Davis et al. 2019) and 6 - 15 X that of *S. cerevisiae* (Landry et al. 2007; Hodgins-Davis et al. 2019). However, it is difficult to directly compare *V_m_*between different organisms due to numerous factors including varying experimental designs, inclusion of all genes versus only DEGs, and technological differences between microarray and RNASeq. Even including these estimates, it is very clear from the within and between species variation in *V_m_* that there is a compelling reason for similar studies in a much broader array of organisms.

Our estimates of *V_m_* also permit calculation of mutational heritability (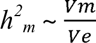, where *V_e_* is the environmental variance) which is useful in comparing how much variation mutation can generate across different traits. We observed a median mutational heritability of per gene expression as 1.4 x 10^-3^ across the 28 MA lines from two strains. A recent review by Conradsen et al (2022) showed that *h^2^_m_* ranged from 2.5 x 10^-5^ to 1.0 x 10^-2^ across numerous traits and taxa. Therefore, our estimate of *h^2^_m_* for gene expression fell in the higher end of this range, but is broadly comparable to the amount of variation generated by mutation in other largely polygenic traits. Notably, microbes and transcriptomic traits were excluded from the review, but the fact that the *h^2^_m_* of gene expression in *C. reinhardtii* is similar to that of other traits implies that mutation can create a substantial amount of variation for natural selection to act upon. As *C. reinhardtii* is haploid, the effects of mutations can not be masked by heterozygosity, which may contribute to higher estimates of *h^2^_m_*. One might assume that the mutational target size of a single gene’s expression is relatively low compared to complex life history traits or fitness, but if there are numerous *trans*-acting effects caused by regulatory network feedback this may not be the case. The *P_TDH3_*-YFP reporter gene in yeast was estimated to have a *trans* mutational target size of 118 Kbp (Metzger et al. 2016). Assuming the *C. reinhardtii* single base mutation rate of 1.2 x 10^-9^ and 900 generations (Ness et al. 2015), this target size implies that 1 in 7.7 genes should experience a regulatory mutation during the MA experiment, and some of these mutations may affect multiple genes.

Mutational variance is often scaled per generation, partly to facilitate estimates of trait evolution over time, but also because the number of mutations causing the change in each line is often unknown. As mutations are the cause of expression change and the expected number of mutations per generation varies between individuals and across species, we may gain insight from per-mutation estimates of mutational variance. Here, we were able to use mutation counts for each line to estimate expression change per mutation (*V_mm_*) using generalized mixed-effect linear modelling. Across genes, we found a 4.917 difference in median *V_mm_* of the strains. These values are both substantially higher than the *V_m_* per generation, which is predicted given that we only expect ∼1 mutation per 10 generations in *C. reinhardtii*. However, while the direction of change between mutation variance scaled per generation and per mutation is expected, the magnitude is much larger than expected based on the number of mutations per generation (*V_mm_* is 24317 *V_m_*). This discrepancy may in part be due to the evolution of mutator strains among the MA lines, with 3.517 - 817 higher mutation rates than other MA lines from the same strain.

When we excluded these two mutators from our model, *V_mm_* estimates dropped by more than half (CC-2344 *V_mm_* = 0.15; CC-2931 *V_mm_* = 0.028), but not enough to explain the discrepancy between *V_m_* and *V_mm_*. Taken together, the lack of congruence between the mutation rate of the two strains and the amount of mutation variance for expression, as well as the relatively larger *V_mm_* to *V_m_* estimates suggests the relationship between mutation count and the extent of expression divergence is complex, which we explore more below.

### Distribution of expression effects of mutation

Contrary to the simple prediction that more mutations would introduce greater changes in expression, we found no linear correlation between total absolute log_2_-fold expression change and the number of mutations in a MA line (Fig S1, *R^2^* = 0.058, *p* = 0.421). In fact, some MA lines with very few mutations (CC-2344) had the most DEGs and total change in expression (Fig 1). To investigate whether there was a bias in the direction of expression change, we separated genes into up- and down-regulated categories.

Lines that experienced large expression changes tended to accumulate a similar amount of expression change in both directions (Pearson’s *R* = 0.826, *p* = 6.03 x 10^-8^). To ensure that the pattern between mutation count and expression change was not driven by large fold-changes in genes with ancestrally low expression, we also considered the relationship between mutation and read count differences (MA - ancestral read count) but similarly found no correlation (Fig S2).

One explanation for the weak linear correlation that we see between expression change and mutation count is that the underlying distribution of expression effects of mutation (DEE) includes a large number of mutations with little or no effect and a long tail of rare mutations that drive large changes in expression. We therefore attempted to infer the DEE as a gamma distribution that would describe the number of DEGs seen in each of the 28 MA lines as a function of the number of mutations they each harbour. We fit three models, one gamma distribution and two shifted gamma’s with a preset fraction of mutations that introduced 0 DEGs or 0 and 1 DEG. Of the three models, the best fitting distribution was strongly right skewed with 95% of mutations causing 0-1 DEGs. According to this model with a discrete bin of zero-effect mutations plus a gamma, 42% of mutations caused no change in expression, 53% caused 1 DEG and 5% caused 2 or more genes to alter in expression (Fig 2). The long tail of high effect mutations meant that the overall mean number of DEGs per mutation is 14.3. Although we likely do not have enough data to precisely infer the shape of the DEE for mutations of large effect (those causing 2 or more DEGs), the three models reach a consensus. Many mutations have no effect on expression, a fraction of mutations affect a single gene, while the long tail of large effect mutations have the potential to create many DEGs. This model fits with what we know about the effects of *cis* and *trans* regulatory mutations with *cis* mutations affecting only local genes, while *trans*-acting mutations have the potential to alter the expression of many genes through pleiotropy (Wittkopp 2023). The existence of a small fraction of large effect mutation also potentially explains why some MA lines with very few mutations have very divergent transcriptomes while others with more mutations have relatively few DEGs. In fact, when we simulate the expected number of DEGs seen in 28 MA lines with the observed mutation counts, the predicted number of DEGs per line is very similar to observed data (1321 DEGs/MA line predicted, 1394 DEGs/MA line observed).

**Fig 2.**
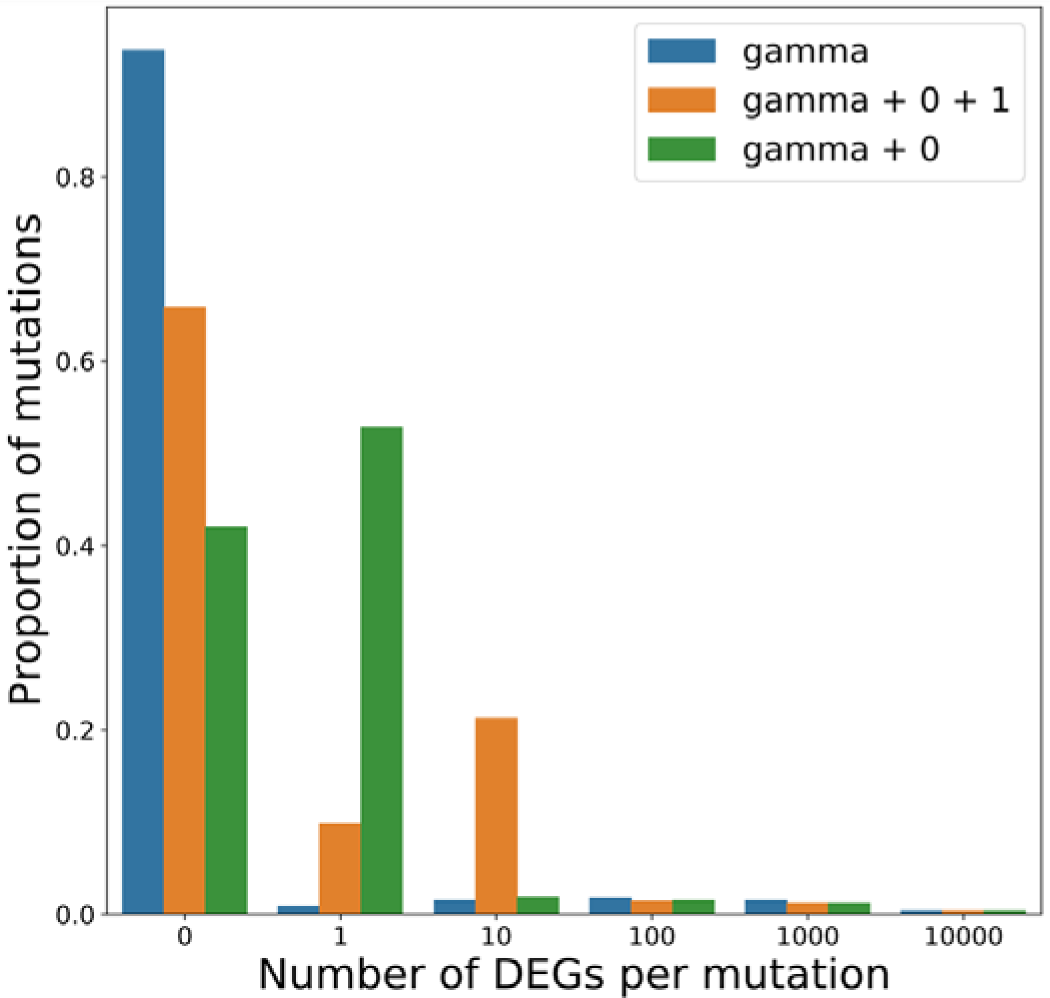
Distribution of expression effects of *de novo* mutations. The inferred distributions describe the number of differentially expressed genes caused by a randomly selected *de novo* mutation. Three distributions were used to fit the mutation and DEG counts of all 28 MA lines. Of the three distributions, the best fitting model (green) included a category of mutations that cause no differential expression (0 DEG) and a gamma distribution shifted by one. The mean number of DEGs per mutation was 14.3 but 95% of mutations were estimated to change the expression of zero or one gene, while the remaining 5% caused 2 or more (0-DEG category =0.42, gamma shape = 0.0118, gamma scale = 2005)

An alternative explanation to our modelled DEE is that the expression of many genes are affected by more than one mutation. Each gene’s expression is the sum of multiple up- and down-regulating mutations creating the possibility of compensatory mutations that would mask the individual effects of mutations. However, under this compensatory model there should be an increase in both the mean and standard deviation of expression change with increasing mutations per line - which is not what we observed.

### Correlates of expression change

It is often hypothesized that genes with high expression are more important to fitness and therefore subject to stronger selective constraints. Indeed, in numerous species there is a negative correlation between expression level and the rate of protein sequence evolution (Pál et al. 2001; Pál et al. 2003; Bloom and Adami 2004; Marais et al. 2004; Rocha and Danchin 2004; Zhang and Li 2004). We find a similar pattern in *C. reinhardtii* with high expression genes displaying reduced sequence divergence relative to *C. incerta* (0.12 for K_a_/K_s high expression genes_ < 0.33 for K_a_/K_s low expression genes_, Brunner Munzel test, *p* = 5.41 x 10^-104^). It is also reasonable to posit that genes with high expression may be critical to the transcriptome and may have evolved robust expression regulation, making them less susceptible to mutational or environmental perturbation. Contrary to this prediction we see relatively little evidence that ancestral expression level predicts the extent of expression change due to mutation (Fig S3). Genes with higher ancestral expression in fact show a slight elevation in fold expression change from the ancestor.

On the other hand, we find a clear correlation that higher expression genes have higher *V_m_* and and *h^2^_m_*; but this reflects the expected mean-variance correlation when *V_m_* is estimated from read counts, such that genes with more reads have more scope to vary in absolute terms. Relatively few other studies have examined the relationship of expression level and susceptibility to mutation. Similar to our findings, there was an overall weak positive relationship of *h^2^_m_* and expression observed in *D. melanogaster* (Fig S4), but that was slightly negative when developmental stages were separated (Rifkin et al. 2005). So while our results support the finding that protein evolution is slower in high expression genes, we find no evidence that the mutational effects are buffered in these high expression genes.

In addition to expression level, the interactions among genes and their relative regulation can be summarized in interaction networks. The centrality-lethality rule describes the concept that highly connected genes in an interaction network are more likely to be critical to fitness (Jeong et al. 2001; Yu et al. 2004; Hahn and Kern 2005; He and Zhang 2006; Yu et al. 2007; Zotenko et al. 2008). There is clear support for this idea from experimental and comparative genomics (Pál et al. 2001; Rocha and Danchin 2004; Subramanian and Kumar 2004; Zhang and He 2005; Liao et al. 2006; Zhang and Yang 2015).

Extending this idea, it is reasonable that the expression level of central genes is also subject to stronger stabilizing selection, and that mechanisms may have evolved to protect central genes from mutations that alter their expression. Using a reconstruction of *C. reinharditii’s* metabolic network, iCre1355 (Imam et al. 2015), we tested whether highly-connected metabolic genes (*i.e*., degree and betweenness centrality) underwent less expression change due to mutation. Contrary to our prediction, we found that the genes with the highest degree (those with the most connections to other genes) had a 25% higher median absolute-fold expression change than the lowest degree genes (low degree genes = 0.35, high degree genes = 0.425, Brunner Munzel test, *p <* 0.03). We also noted that the subset of genes included in the metabolic network had higher expression than the genome average and therefore tended to have higher *V_m_*because of the same mean-variance correlation of *V_m_* and expression noted above. When we look for altered patterns of expression change in the regulatory network, we find a similarly significant positive correlation of connectivity (*i.e.,* intramodular connectivity) in a regulatory module with the median expression change of a gene although the amount of variation explained is relatively low (*R*^2^ = 0.04, *p* = 0.001). The positive relationship between connectivity and expression divergence does not support the idea that mechanisms have evolved to buffer these genes from mutational effects, but nor does it rule it out. Given the interconnectedness of hub genes, it may be expected that they would be more prone to perturbation and the fact that their mutation vulnerability is quite similar to other genes could be a consequence of these buffering mechanisms. No previous mutation accumulation studies have examined the relationship between network topology and mutational effects on expression. However, in *D. melanogaster* (Lemos et al. 2004; Lemos et al. 2005) reduced divergence and polymorphism of expression level was seen in highly connected proteins, and if this trend holds in *Chlamydomonas* it would imply that there is stronger stabilizing selection removing the incoming variation from mutation.

### Evidence of cis- and trans-acting mutations

The inferred DEE suggested a large fraction of mutations that cause only 1 or a small number of DEGs, which could represent mutations in *cis*-acting regulatory regions that affect local genes. We therefore sought to test for an over-representation of mutations near DEGs, under the simplifying assumption that *cis*-mutations are near the genes they affect. To generate a null distribution of the expected distance of a DEG to its nearest mutation, we conducted 10,000 iterations where the relative positions of mutations and DEGs were randomized. To reflect the non-uniform distribution of mutations, we retained their position in the genome and randomly assigned genes as DEGs. When we compared the distribution created with observed DEGs to the null, we found an enrichment of DEGs with mutations within 100 bp of the gene flanks in 12 of 28 MA lines (permutation test, *p* < 0.05; Fig S5) and an additional six lines with marginally significant *p* values (permutation test, 0.05 < *p* < 0.1). In Fig 3, we display a representative sample (CC-2344 L14) showing evidence of an over-abundance of mutations near DEGs (red) relative to the expectation if there was no relationship (blue). We also found several mutations within DEGs, which were twice as likely to be found in UTRs as would be expected based on the proportion of UTR sequences in the genome. This may be expected given that UTRs are enriched with promoters/enhancers and are known to regulate RNA transcription, stability, and translation (Steri et al. 2018). We also noticed that MA lines with many DEGs were less likely to show the enrichment of nearby mutations. This may be due to the presence of one or more *trans*-acting mutations with multiple regulatory targets that masked the signals of any *cis*-acting mutations that only affect one gene. Though previous experimental results suggest that *cis*-acting mutations tend to down-regulate genes (Metzger et al. 2016), we did not see a bias in the direction of expression change in DEGs that were <100 bp from a mutation. From single gene reporter systems (Metzger et al. 2016), we have estimates that the *trans*-acting mutation target is 385-fold larger than the *cis*-target for that single reporter gene. However, if *cis* acting mutations are gene-specific, and *trans*-acting mutations are pleiotropic, then the full DEE of all genes might have a larger overall proportion of target sites for *cis*-acting mutations and could reconcile why we see a relatively clear signal of mutations near genes.

**Fig 3.**
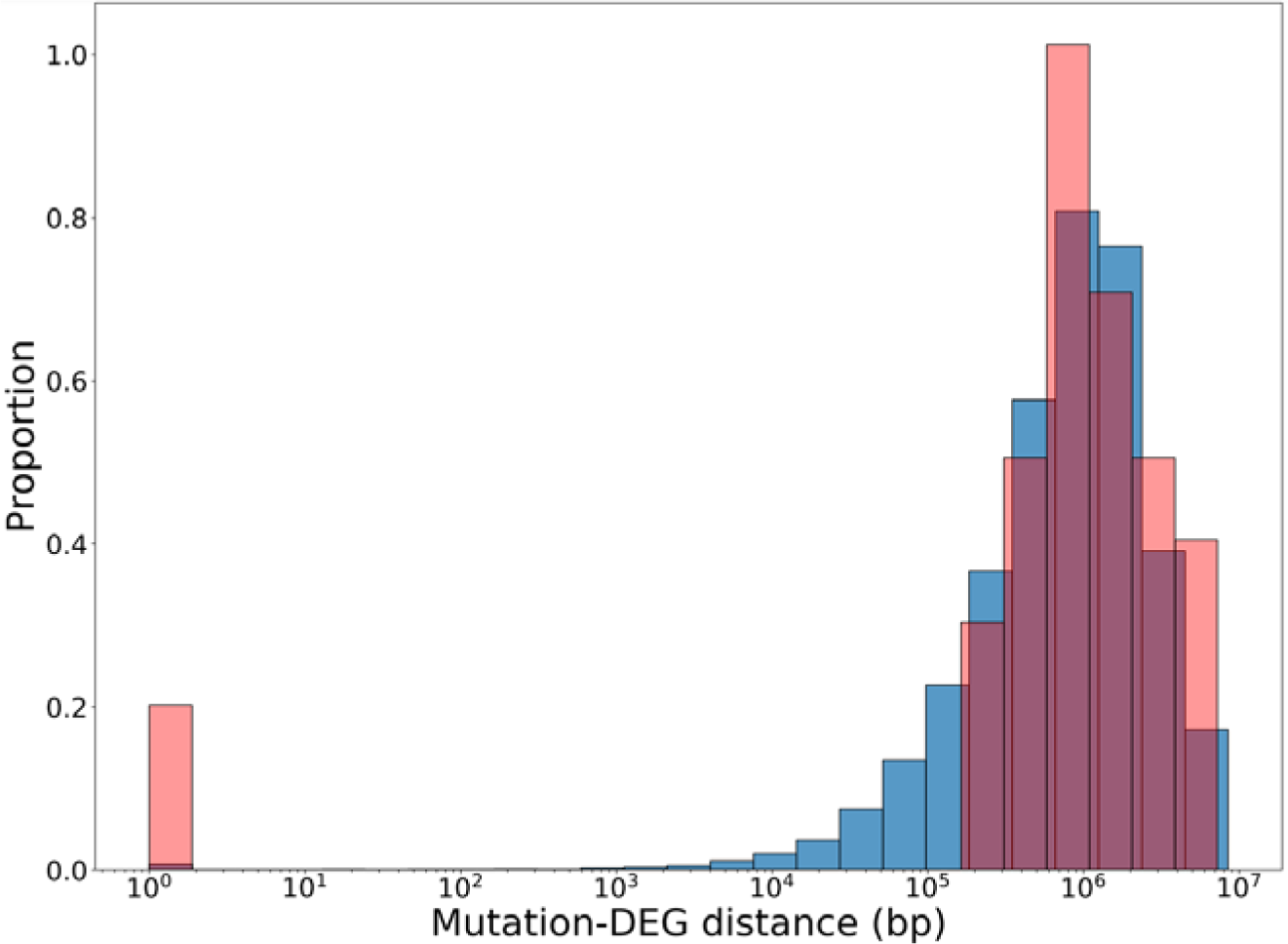
Evidence of *cis*-regulatory mutations. Distribution of distance between a DEG and its nearest mutation for observed (red) and simulated data (blue). The observed distribution is one representative MA line and the expected distribution is based on 10,000 trials where the positions of DEGs in the genome are randomized. The excess of DEGs co-localizing with mutations is apparent in the excess of observed data on the left of the distribution. Mutations found within the coding or non-coding regions (UTRs and introns) were assigned a distance of 1.

Although the majority of the inferred DEE is small effect mutations, the long tail of mutations causing multiple DEGs may indicate the presence of *trans*-acting sites with pleiotropic effects. It is clear from the ratio of DEGs to mutations (mean 15.7 DEGs/mutation) that there are many mutations that affect multiple genes, especially given that many mutations will have little or no effect on expression. We also found that DEGs tended to be clustered near one another in the *Chlamydomonas* co-expression network (Strenkert et al. 2019). The shortest path between pairs of DEGs from the same MA line was less than the shortest path length between random pairs of genes in 14 out of 28 MA lines (Permutations test, *p* < 0.05). Fig 4 is a representative plot of one MA line (CC-2344 L1), where the signal of co-expression network clustering can be observed as an excess of short paths between DEGs. A similar pattern was seen in *C. elegans* where gene expression changes from mutation accumulation clustered in the network, and were interpreted as evidence of *trans*-acting mutations. It has been shown in *S. cerevisiae* that the topology of the co-expression network can explain the pleiotropic effects of *trans*-acting mutations, in line with what we observe. Significant clustering of DEGs could be the consequence of mutation in a common *trans*-acting factor that controls expression in a given co-expression module or regulatory feed backs that result from alterations to the expression or structure of related genes.

**Fig 4.**
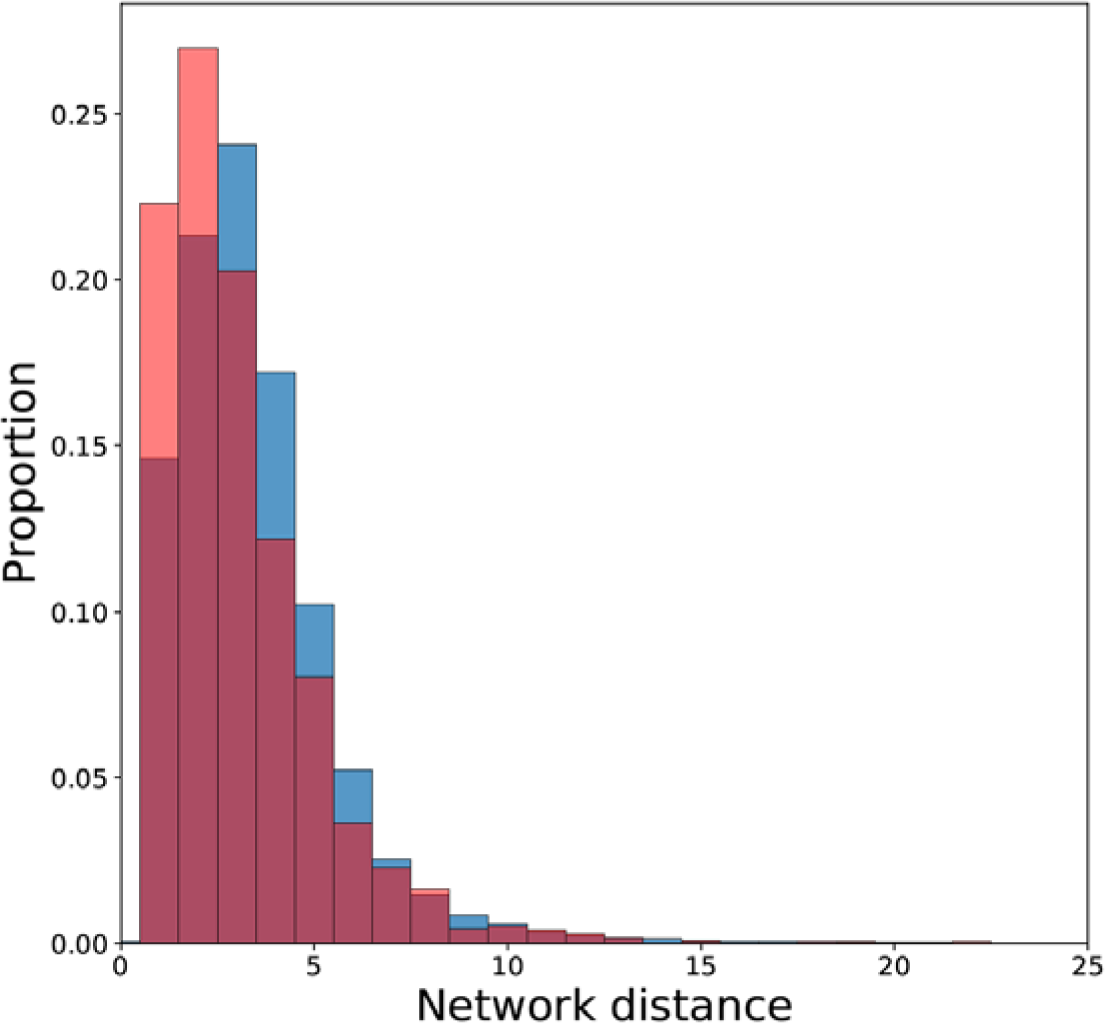
Clustering of DEGs in the co-expression network. Overlaid distributions of the shortest path lengths between observed DEGs (red) and the null (blue) expectation. The null distribution was created by calculating the shortest paths among a random sample of genes from the co-expression network. The figure above is a representative plot of a single MA line demonstrating an excess of observed short paths among DEGs at the left of the plot indicative of clustering (Permutations test, p < 0.05). 14 of 28 MA lines had a significant excess of clustered DEGs.

### Fitness consequences of expression change

It has been shown that expression is often under stabilizing selection (Denver et al. 2005; Rifkin et al. 2005; Huang et al. 2016), thus we would expect MA lines with more total expression change to equally experience more fitness decline. Consistent with this prediction, we found a negative correlation between the number of DEGs per MA line and fitness (Fig 5, *R*^2^ = 0.233, *p* = 0.032), and a similar relationship between the sum of absolute log_2_-fold change and fitness (Fig S6; *R*^2^ = 0.2, *p* = 0.034). It is worth noting that the correlation was dependent on a single influential data point with high expression divergence and low fitness at the bottom right of Fig 5 (excluding this data point, *R*^2^ = 0.183, *p* = 0.201). However, there is nothing otherwise unusual about this particular data that would justify its exclusion, so while the relationship is not strong, more replication would be required to determine if the relationship between fitness and expression is real. The association between declining fitness and increasing number of DEGs implies the deleterious nature of large changes in the transcriptome due to mutation. Zande et al (2022) had similar findings in yeast, where *trans* mutations had negative fitness effects due to pleiotropy. It is also worth noting that these same lines have previously been analyzed, where it was seen that there was a negative relationship between the number of mutations and fitness (Kraemer et al. 2017; Böndel et al. 2019; Böndel et al. 2022), but given that we did not seen a relationship between the amount of expression divergence and mutation count the contribution of some deleterious mutations may have been to non-expression related traits while others influenced expression to cause an overall decline in fitness. In combination with our results which describe the effects of a random set of *de novo* mutations, evidence suggests that *C. reinhardtii* is vulnerable to harmful mutations that disrupt the regulation of gene expression.

**Fig 5.**
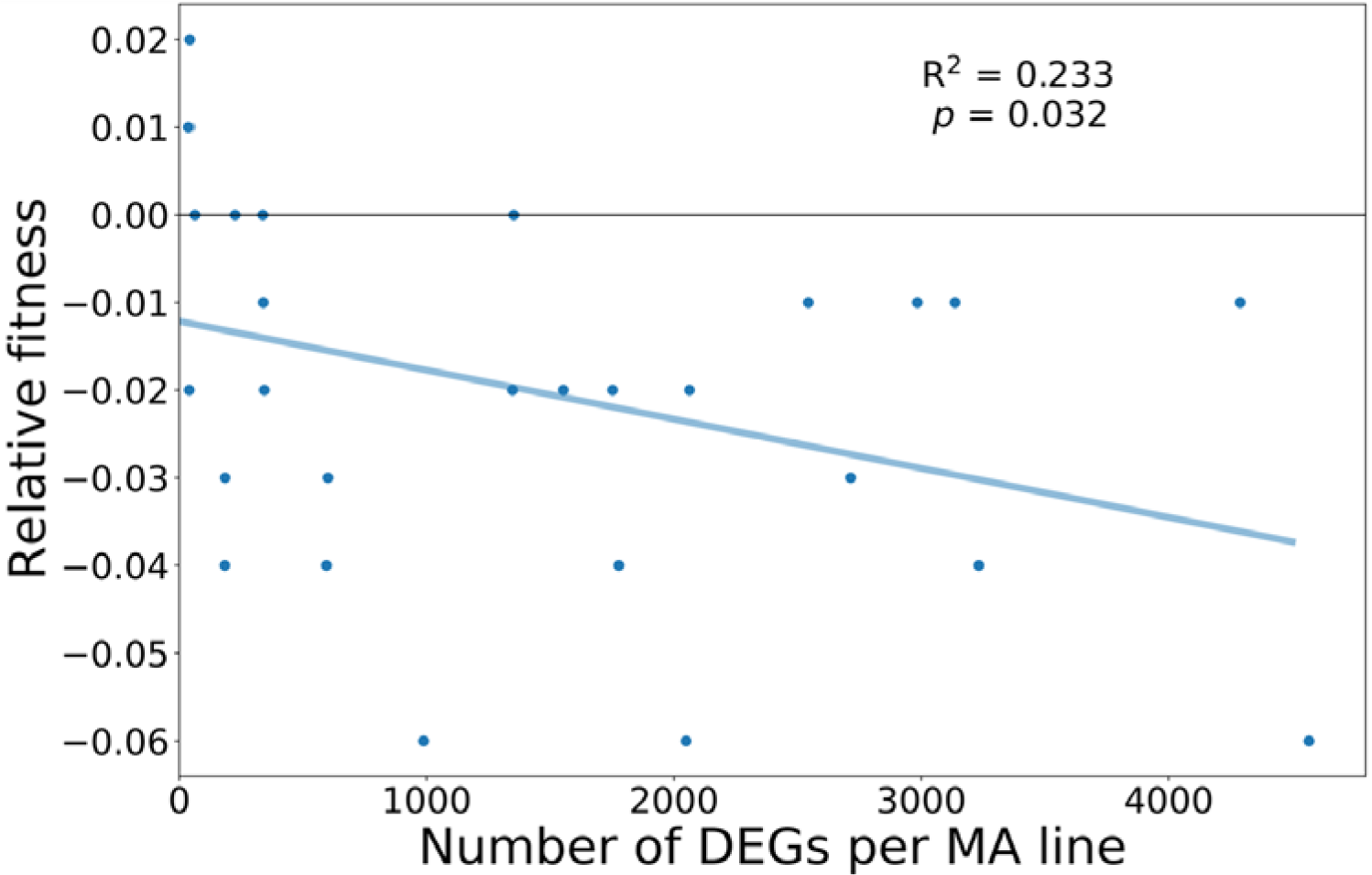
Evidence of fitness decline in lines with more differentially expressed genes. Linear regression modelling the negative relationship between the number of significant differentially expressed genes in each MA line and its fitness relative to the unmutated ancestor (*R^2^* = 0.233, *p* = 0.032).

Taken together we see clear evidence that mutation can generate a substantial amount of genetic variation in gene expression and this new variation is important for organismal performance. For example, it has been a long-held belief that the contribution of mutation to genetic variance is less than 1% of the environmental variance. But, we found that mutation may contribute significantly more to variation per generation than formerly considered, almost 24%. This is a considerable amount of mutational variation that is heritable, which may change the perception of the raw materials available for evolution via selection to act upon. Using mutation accumulation lines allows us to observe the cumulative effect of mutations from the spectrum of naturally occurring mutations. Our estimated DEE indicates that there is a substantial mutational target for large effect mutations that pleiotropically generate expression changes across the transcriptome. There are some lines harbouring tens of mutations showing thousands of differentially expressed genes. It seems likely that these pleiotropic mutations are driven not only by mutations in *trans*-acting regulators like transcription factors, but also by alterations in the balance of the metabolism of the cell and feedbacks in the regulatory network. Despite this, we see no evidence for mutational robustness in genes central to the metabolic and co-expression network. A limitation of our data is that we can not link the expression changes to individual mutations and therefore can not assess the nature of these large effect mutations. Therefore, the possibility of compensatory mutations means that some expression change may be masked by subsequent mutations. With current technology it is not cost-effective to map *trans*-acting mutations to their expression effects but advances in single cell technologies may allow for such resolution in the near future.

## Acknowledgments

We would like to thank Dr. Peter Keightley and anonymous reviewers for helpful comments on our paper. We would also like to thank Dr. Jarrod Hadfield for invaluable help in estimating mutational variance. This work was supported by a Natural Sciences and Engineering Research Council (NSERC) Discovery grant (RGPIN/06331-2016), the Canadian Foundation for Innovation John R. Evans Leaders fund (35591) to R.W.N. and an NSERC graduate scholarship to EJB.

## Supplementary materials

**Fig S1:**
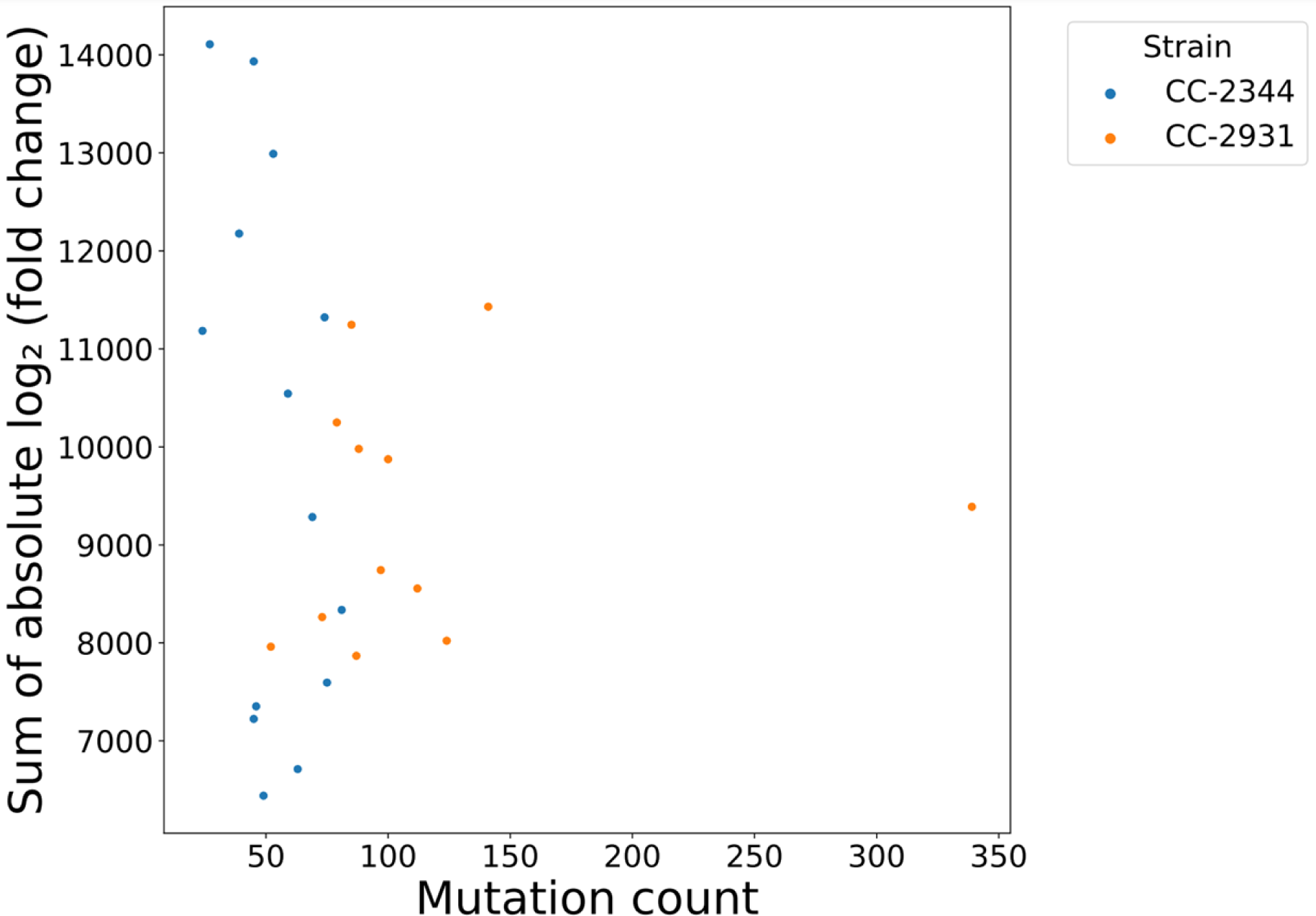
Scatterplot of the sum of the absolute log_2_ fold change by mutation count. Expression change is measured as the sum of absolute log_2_ fold change across all genes in an MA line and is plotted against the number of mutations in that MA line. There was no significant correlation found between the two variables *(R^2^* = 0.058, *p* = 0.421).

**Fig S2.**
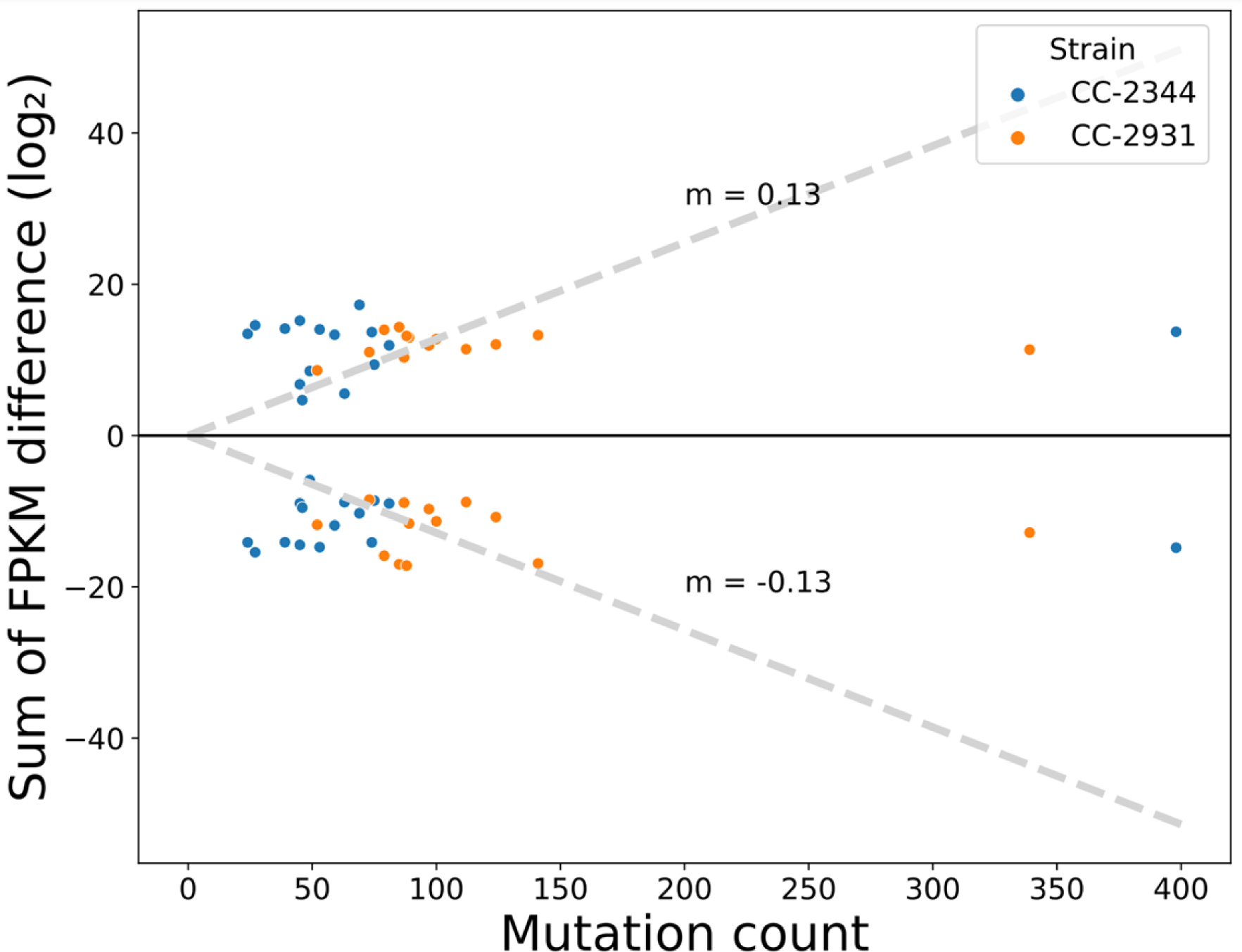
No correlation between the sum of FPKM differences and mutation count. Plot of the log_2_ transformed summed FPKM-normalized read count differences between MA lines and their respective unmutated ancestor. Only differentially expressed genes with *p*_adj_ < 0.05 were considered. Each MA line is shown twice with the same mutation count, points above the y=0 line represent upregulated genes and below the line are down-regulated genes. The dashed line represents the average rate at which expression changes per mutation. We did not find a correlation between the non-transformed differences in expression and mutation count (positive expression change ∼ Pearson *R* = −0.0825, *p* = 0.676; negative expression change ∼ Pearson *R* = −0.0607, *p* = 0.759).

**Fig S3.**
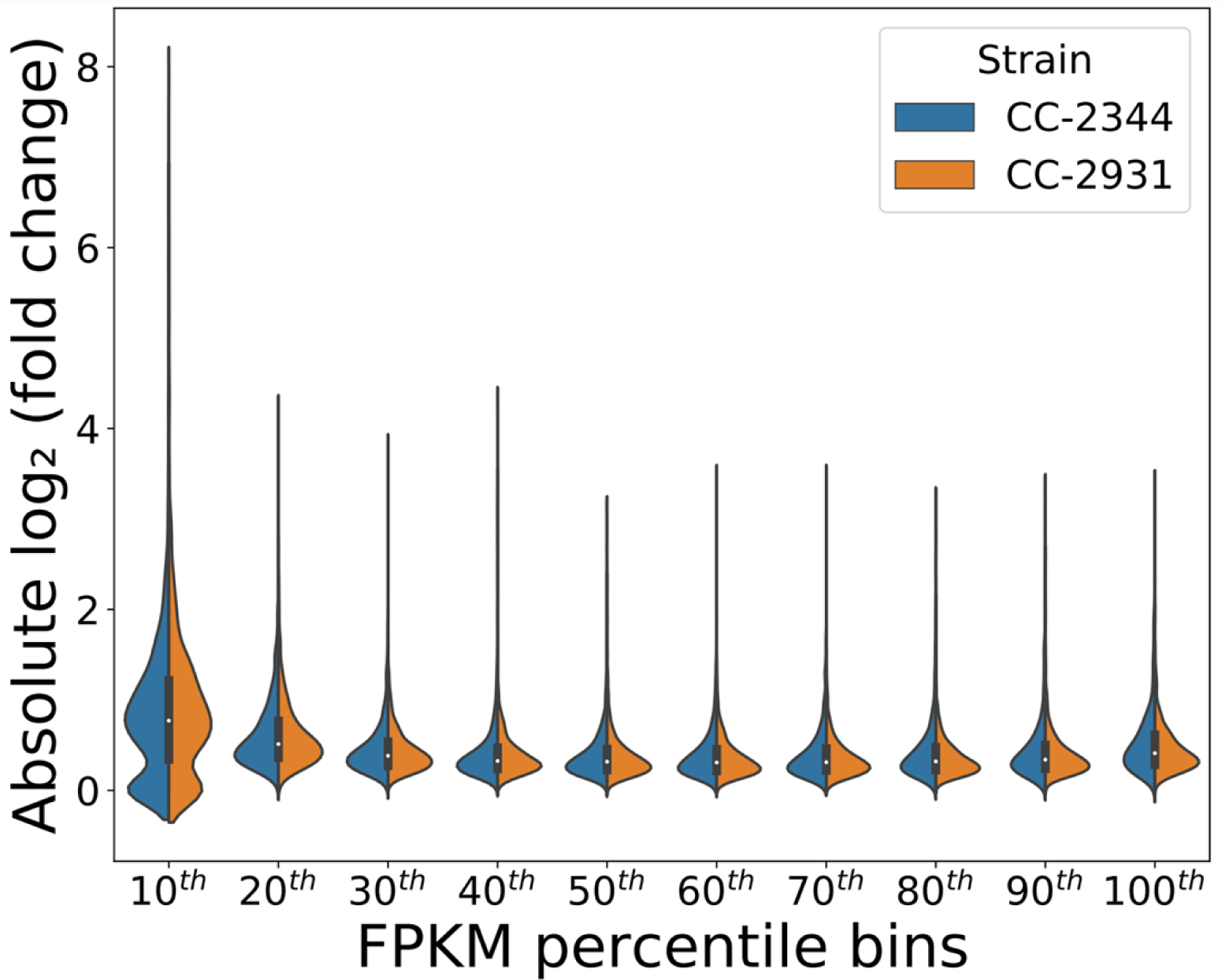
Ancestral expression level does not predict absolute expression change. Split-violin plot showing the distribution of genes in each FPKM-normalized ancestral expression bin (n_CC-2344_ = 1745, n_CC-2931_ = 1737). Each measurement within the distribution represents the median absolute expression change across all 28 MA lines per gene. There is no relationship between the ancestral expression level of a gene and its respective degree of expression change. The median expression change decreases from the 10^th^ to 50/60^th^ percentile expression bin and increases thereafter.

**Fig S4.**
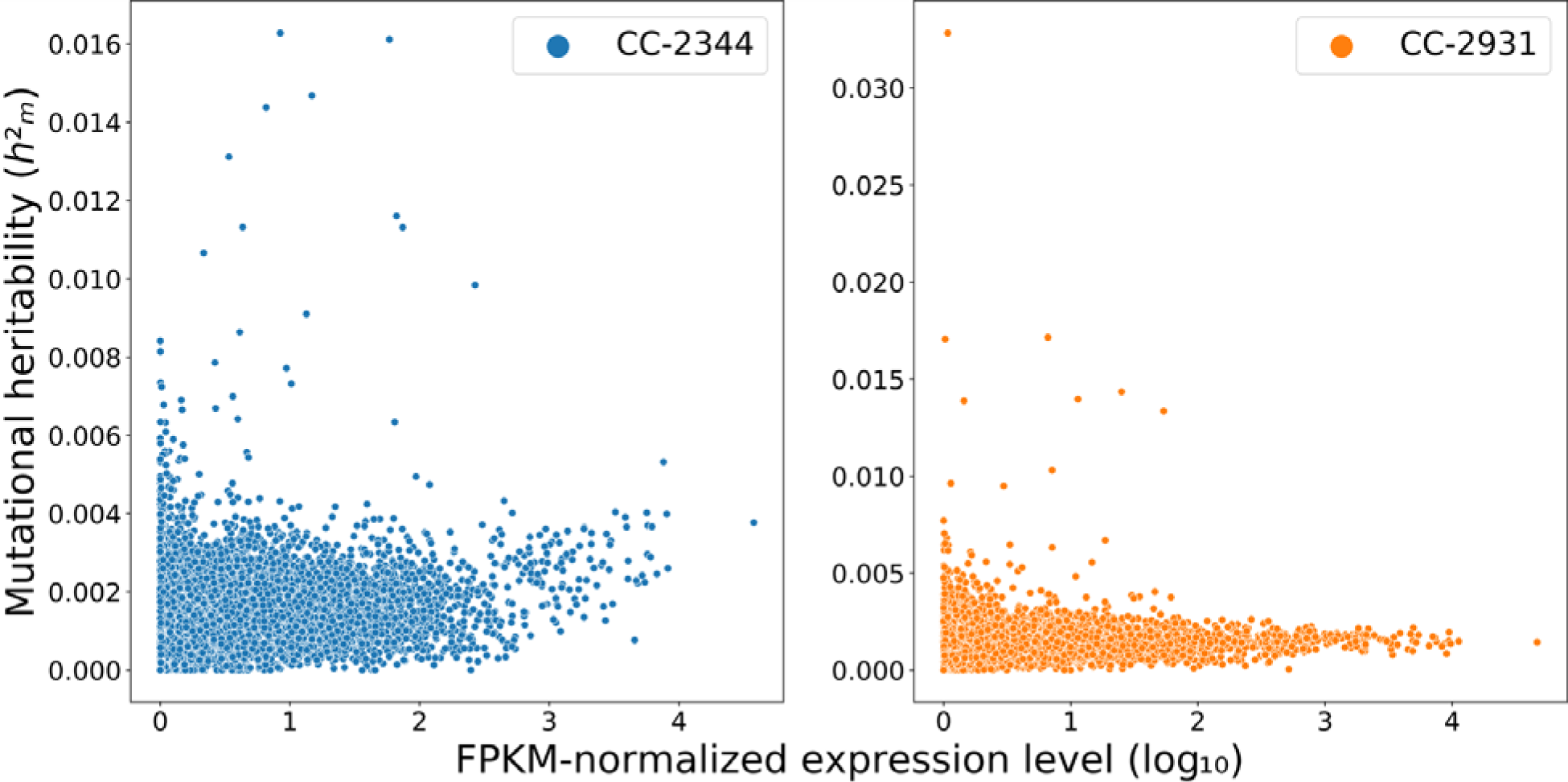
Slight positive correlation between mutational heritability and expression level. Scatterplot of the FPKM normalized gene expression counts of the ancestral line of each strain against its respective mutational heritability. The mutational heritability was estimated using the per generation mutational variance. There is a weak positive relationship for CC-2344 (Pearson’s *R* = 0.20, *p =* 1.22 x 10^-162^) and CC-2931 (Pearson’s *R* = 0.06, *p =* 7.59 x 10^-14^).

**Fig S5.**
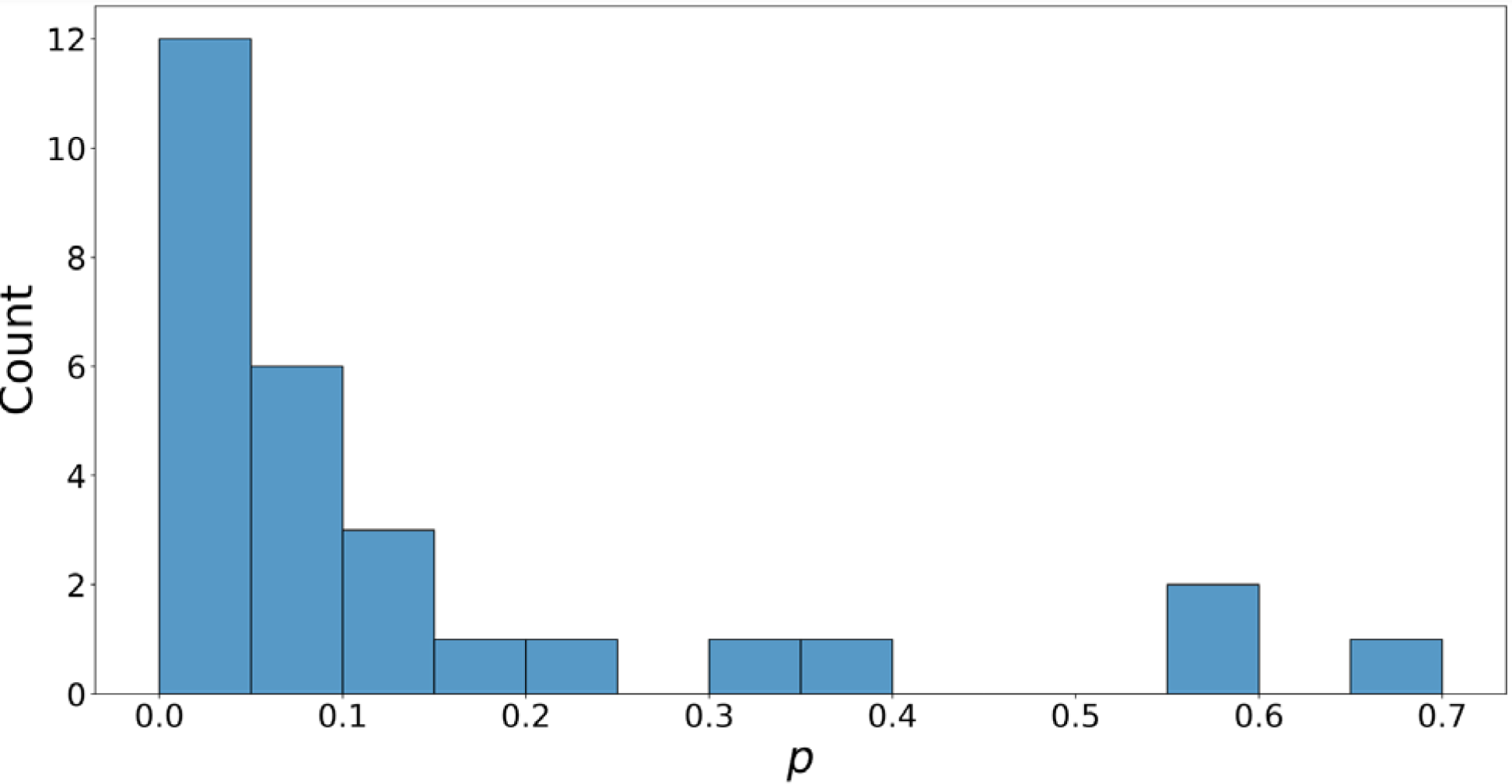
Overrepresentation of mutations within 100 bp of differentially expressed genes. Evidence of *cis*-regulatory mutations was inferred from an over-representation of mutations within 100 bp of differentially expressed genes (DEGs). The observed distance between mutations and their nearest DEGs was compared to the expected distribution created by randomizing the DEGs across the genome over 10000 iterations. The *p* values represent the fraction of trials where the cases of simulated DEGs co-localizing with mutations was greater than those with observed DEGs. The distribution of *p* values is heavily right-skewed, showing the presence of an excess of mutations in or near DEGs in most MA lines.

**Fig S6.**
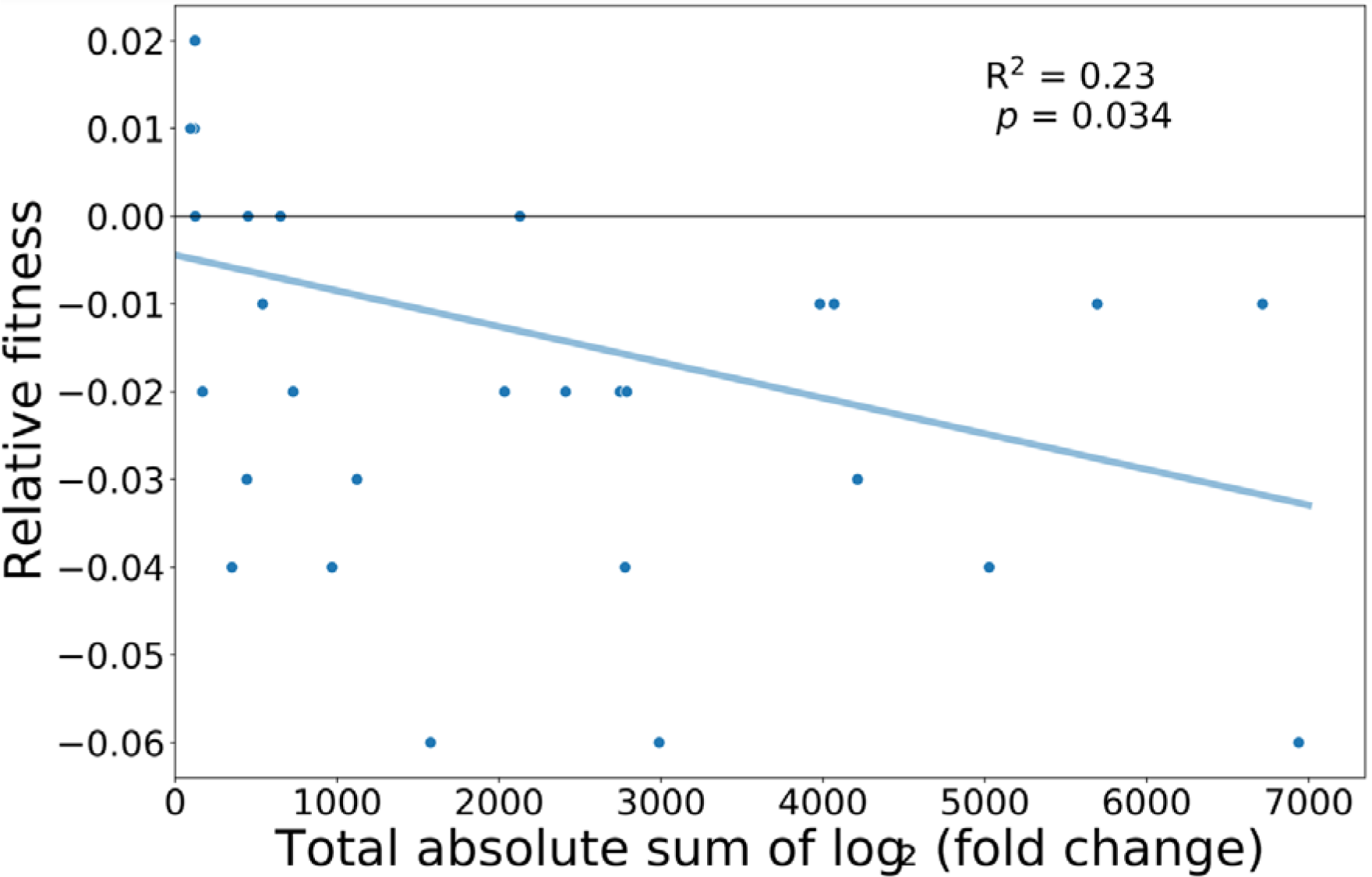
A negative correlation between relative fitness and total absolute sum of log_2_ fold change. Linear regression modelling the relationship between the total sum of absolute log_2_ fold change in differentially expressed genes and its fitness relative to the unmutated ancestor (*R^2^*= 0.23, *p* = 0.034).

